# DATS: the data tag suite to enable discoverability of datasets

**DOI:** 10.1101/103143

**Authors:** Susanna-Assunta Sansone, Alejandra Gonzalez-Beltran, Philippe Rocca-Serra, George Alter, Jeffrey S. Grethe, Hua Xu, Ian Fore, Jared Lyle, Anupama E. Gururaj, Xiaoling Chen, Hyeon-eui Kim, Nansu Zong, Yueling Li, Ruiling Liu, Burak Ozyurt, Lucila Ohno-Machado, bioCADDIE Working Groups Members

## Abstract

Today’s science increasingly requires effective ways to find and access existing datasets that are distributed across a range of repositories. For researchers in the life sciences, discoverability of datasets may soon become as essential as identifying the latest publications via PubMed. Through an international collaborative effort funded by the National Institutes of Health (NIH)’s Big Data to Knowledge (BD2K) initiative, we have designed and implemented the DAta Tag Suite (DATS) model to support the DataMed data discovery index. DataMed’s goal is to be for data what PubMed has been for the scientific literature. Akin to the Journal Article Tag Suite (JATS) used in PubMed, the DATS model enables submission of metadata on datasets to DataMed. DATS has a core set of elements, which are generic and applicable to any type of datasets, and an extended set that can accommodate more specialized data types. DATS is a platform-independent model also available as a Schema.org annotated serialization to be used beyond DataMed, for example, in projects like DataCite.

## Introduction

The biomedical and healthCAre Data Discovery Index Ecosystem (bioCADDIE, https://biocaddie.org) is an international joint effort of researchers, informaticians, data scientists, IT professionals, governmental agencies, services providers, publishers and the NIH to facilitate the discovery of available biomedical datasets that are spread across different databases, repositories and on the cloud, through the development of a Data Discovery Index (DDI). The NIH BD2K^1^ DDI Consortium is a direct response to the key recommendations made in 2012 by the Data and Informatics Working Group (https://acd.od.nih.gov/diwg.htm), set by the Advisory Committee to the Director of the NIH, that requested the development of a catalog of biomedical data. bioCADDIE started effectively when strategies and activities needed to implement the DDI were released as a white paper in 2015^2^.

The DDI prototype, named DataMed (https://datamed.org), was first launched in 2016. Currently, it enables browsing and searching over 50 biomedical data sources, getting recommendations on datasets that best satisfy the users’ specific interests, preferences, and needs. Also the ISA-formatted metadata associated to each Data Descriptor article in *Scientific Data* (http://scientificdata.isa-explorer.org) are being index and will be available in the next DataMed release.

DataMed focuses on Findability (F) and Accessibility (A) of datasets. Along with Interoperability (I) and Reusability(R), F&A compose the four elements of the widely-endorsed FAIR principles^3^, which put a specific emphasis on enhancing the ability of machines to automatically discover and use digital objects, in addition to supporting their reuse by individuals throughout their life cycle. DataMed’s architecture encompasses: (i) a repository ingestion and indexing pipeline, which maps the disparate metadata from the diverse repositories, into the unified DATS model; (ii) a terminology server to ensure metadata consistency at a semantic level; and (iii) a web application based on the search engine, which uses the discovery index for locating the appropriate datasets from the set of repositories, and also the terminology server to expand user queries.

DATS metadata elements are used for indexing and searches in DataMed, and are presented as two modules: a core and extended set of elements. The DATS model has been specifically named to echo the JATS metadata elements and format (https://jats.nlm.nih.gov) required by PubMed (https://www.ncbi.nlm.nih.gov/pubmed) to index publications. Like JATS, the core DATS elements are generic and applicable to any type of dataset. As described in the DataMed paper^5^, the extended DATS includes additional elements. Some of these elements are specific for life, environmental and biomedical science domains and can be further extended as needed. The similarity and complementarity between DataMed and PubMed are not a simple convenience: it reflects the ecosystem of interconnected resources that are necessary to support and enable biomedical research as a digital research enterprise, known as the NIH Commons^4^. As the Data Science culture grows, all digital research outputs (not just publications and datasets, but also software, code, workflows, training materials, tools, standards, etc.) must be FAIR. The goal is to empower scientists to fully utilize existing outputs to accelerate their research.

Throughout the project, the bioCADDIE team has undertaken a variety of community-driven activities to design and deliver DataMed; a detailed description of our journey and the resulting implementation are the focus of a complementary DataMed article. Here we focus on presenting the DATS model, its design principles, the methodologies used, the serializations and implementations to date.

## Results

The DATS model describes the metadata elements and the structure for datasets, and powers DataMed’s ingestion and indexing pipeline, as well as its search functionality. The model also follows the FAIR best practices for data management, and the recently released “Candidate Recommendations on Data on the Web Best Practices” (https://www.w3.org/TR/dwbp) by the Data on the Web Best Practices Working Group (WG) under the World Wide Web Consortium (W3C). Both community practices support that data on the web should be discoverable and understandable by humans and machines; and that such usage should also be discoverable and the efforts of the data publishers recognized.

The work has been driven and executed by the bioCADDIE team with the input and feedback of international researchers, service providers and knowledge representation experts. Community participation was coordinated via the bioCADDIE’s Descriptive Metadata WG3 and the Accessibility Metadata WG7; participants are listed in the Acknowledgement section. The Metadata WG is a joint activity with a wider Metadata WG - encompassing other BD2K centers of excellence - and closely connected to the NIH BD2K Center for Expanded Data Annotation and Retrieval^6^ and ELIXIR activities in Europe (https://www.elixir-europe.org).

In August 2015, the first version (v1.0) of the DATS specification was released^7^, with the model available as machine readable JSON (http://json-schema.org) schemata and with several examples. This initial model and each of the following versions were tested by the DataMed development team, with a variety of data sources, and reviewed by the bioCADDIE’s Descriptive Metadata WG3 members and the larger community. As a result of this iterative process, specifications v1.1^8^ and v2.0^9^ were released in March and June 2016, respectively. The latter version represents a further evolution of the model, including the additional metadata elements produced by the Accessibility Metadata WG7, a set of JSON-schemas and JSON examples, and a Schema.org-annotated context file. The current version v2.1^10^ also includes edits based on feedback from a dedicated bioCADDIE repository workshop (https://biocaddie.org/biocaddie-repository-workshop-june-23-2016) and a Schema.org-annotated JSON-LD (http://json-ld.org) serialization.

The following sections provide a description on the current DATS model v2.1; the specifications, serializations and examples are freely available from the bioCADDIE Github repository (https://github.com/biocaddie/WG3-MetadataSpecifications), where further versions will be released, and enquiries and feedback can be submitted.

### Scope and Coverage: Focus on Discoverability

First and foremost, it is important to clarify that DATS has not been designed to perfectly represent each and every step of an experiment; this level of detail and metadata are left to the specific repositories that DataMed works with to index. The scope here is discoverability: help researchers find and access datasets that are stored across a variety of repositories, by providing enough consistent information to help them find what they need, then point them to where and how to access it. Therefore, DATS has been designed with the objective to identify these key metadata descriptors and relations among them.

Developed iteratively, the model is the result of three complementary approaches, detailed in the Methods section; briefly: (i) a review of existing meta-models, (ii) an analysis of the use cases (top-down), and (iii) a mapping of existing metadata schemas (bottom-up), to find convergences and common metadata elements. Such a combined approach has been necessary to define the appropriate boundaries and level of granularity: which queries will be answered in full by the DataMed prototype, which only partially, and which queries are out of scope.

The DATS model is designed around the *Dataset* element, catering for any unit of information stored by repositories. *Dataset* covers both (i) experimental datasets, which do not change after deposition to the repository, and (ii) datasets in reference knowledge bases, describing dynamic concepts, such as “genes”, whose definition change over time. To implement the concept that DataMed is built as part of an interlinked ecosystem of resources, the *Dataset* element is linked to other digital objects, which are the focus of other indexed resources; specifically *Publication* (e.g. PubMed), *Software* (e.g. BD2K Aztec: https://aztec.bio), *DataRepository* and *DataStandard* (e.g. BioSharing^11^, https://biosharing.org). The latter is especially important in the light of the vast swathes of data that still remain locked in esoteric formats, are described using *ad hoc* or proprietary terminology or lack sufficient contextual information. Knowing if a repository uses open community standards to harmonize the reporting of its different datasets, will provide researchers with some confidence that these datasets are (in principle) more comprehensible and reusable.

Key information about the *Dataset* element is its accessibility, which is represented by the *Access* metadata element that encompasses information on authorization, authentication, and access type. Many types of data in the biomedical domain are restricted to protect confidential information about human subjects. Researchers want to know which data sets are readily available and which ones require prior approval, institutional data use agreements, and security clearances. The *DatasetDistribution* element, linked to the *DataRepository, DataStandard* and *Access,* also informs researchers about which data can be accessed directly by machines through an application programming interface (API).

### Modular Model: Domain-Agnostic Core with Extended Elements

The Google JSON style guide (https://google.github.io/styleguide/jsoncstyleguide.xml) has been used to name relevant elements in the DATS model; the requirement levels follow the standard RFC 2119 (https://tools.ietf.org/html/rfc2119). The descriptors for each metadata element (Entity), include: Property (describing the Entity), Definition (of each Entity and Property), Value(s) (allowed for each Property). In both core and extended DATS, Entities are not mandatory, but applicable only if relevant to the dataset to be described; when an Entity is used, only some of its Properties are defined as mandatory. For example, for the 18 metadata elements of the core there are only a total of 10 mandatory properties. An overview of the core and extended elements, their types and relations are shown in Figures 1 and 2, respectively; Figure 3 provides a view of the core entities highlighting the few properties defined as mandatory. DATS has been designed to specifically drive discoverability: to find and access datasets via key metadata descriptors. DATS is not meant to perfectly model an experiment, which is the scope of many repository-level models and schemas. The DATS model may appear quite detailed in places as a consequence of (i) the combined approaches used to identify the required metadata elements (see section 2), and (ii) the attempt to aim for the maximum coverage of use cases with minimal number of metadata elements. Nevertheless, it is anticipated that not all use cases can be fulfilled, and that it is difficult to foresee all types of data sources the DataMed prototype should retrieve information from.

**Figure 1.**
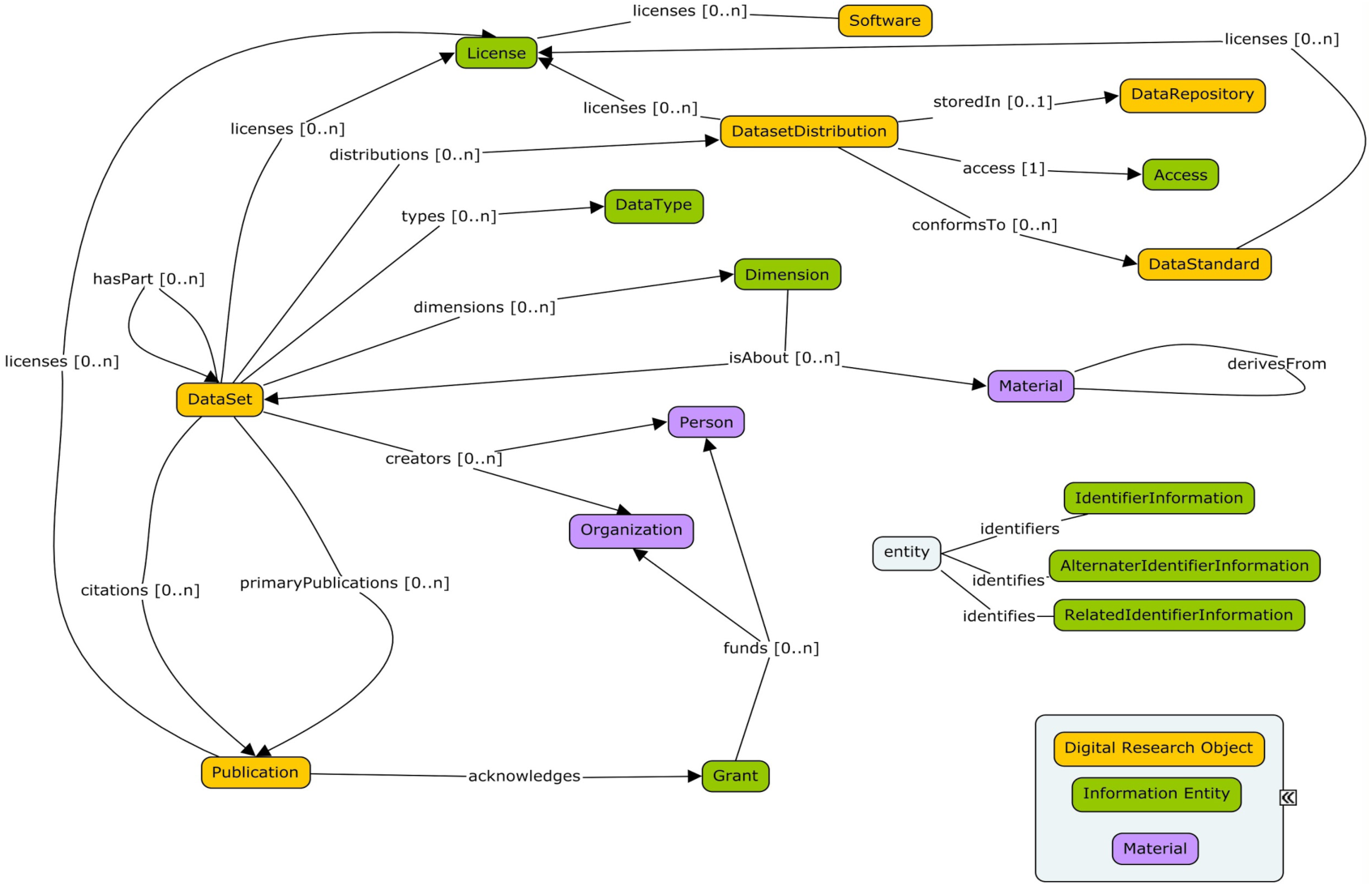
A schematic overview of the DATS core elements, their types and relations.

**Figure 2.**
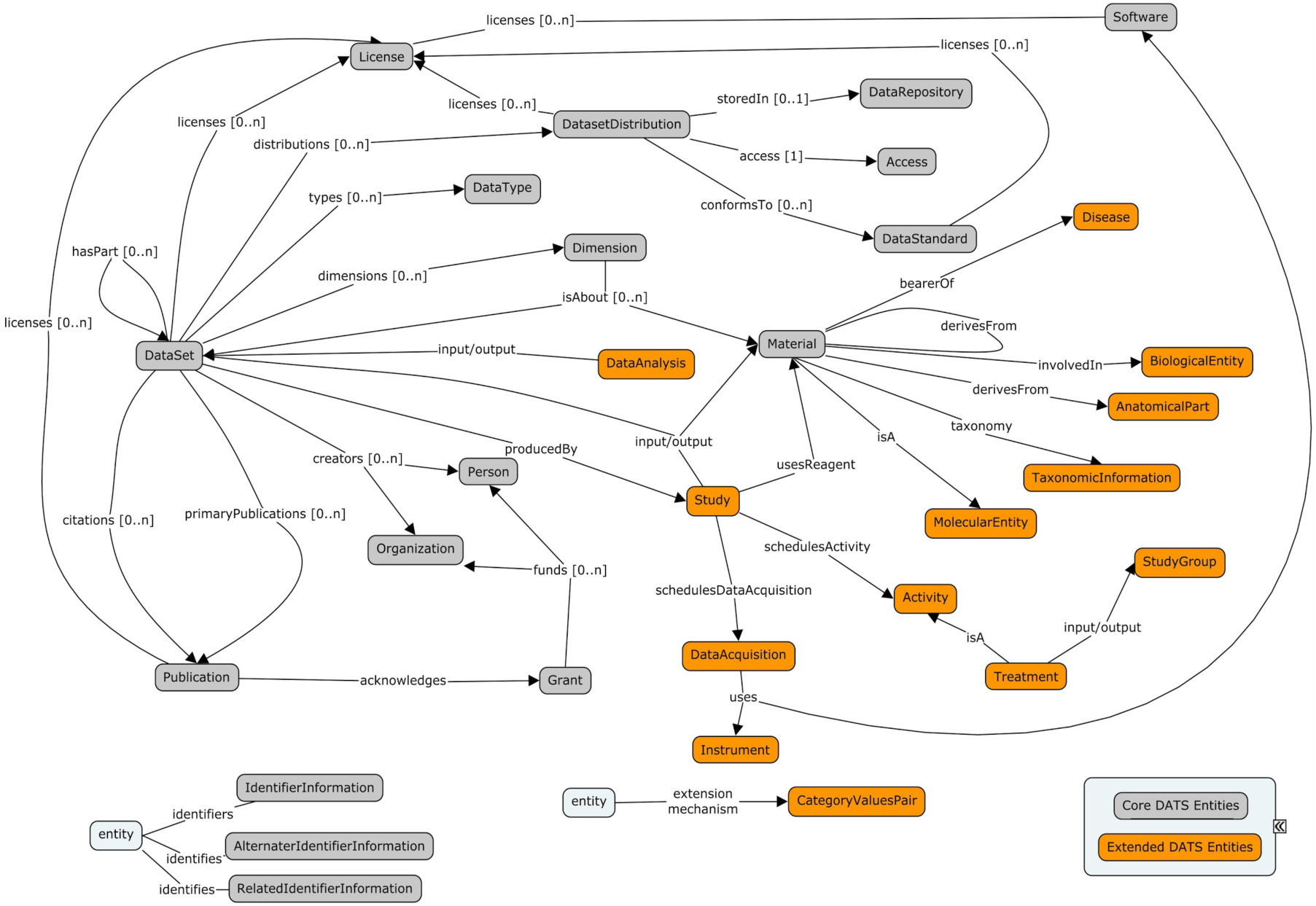
A schematic overview of the DATS core and extended elements, their types and relations.

**Figure 3.**
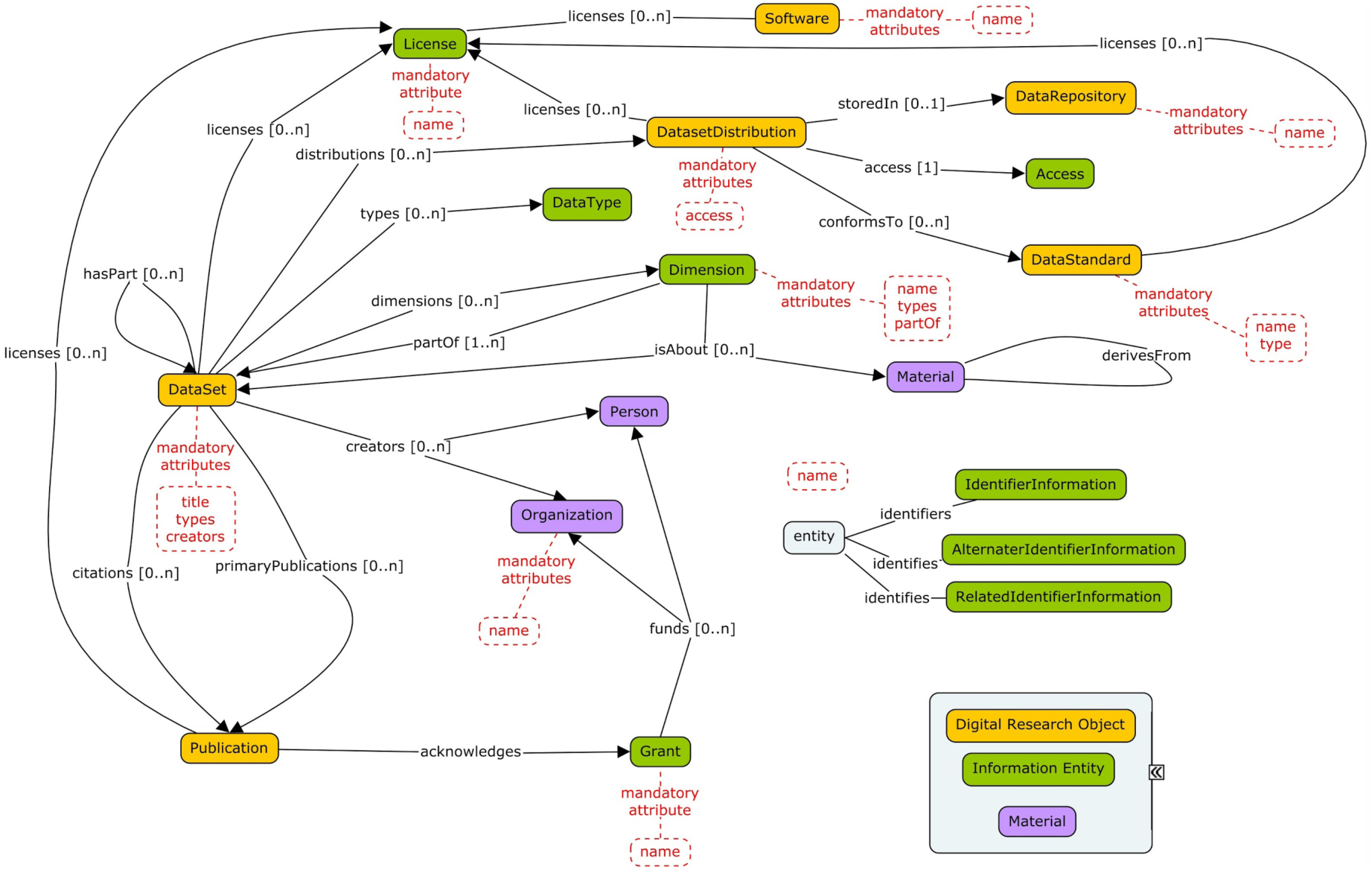
A schematic overview of the DATS core entities and theirs few properties with requirement level “MUST”.

Like the JATS, the DATS core elements are generic and applicable to any type of datasets, and cover basic information, for example: what the dataset is about (using the *Material* entity), how it was produced and which information it holds (*DataType*, *Information*, *Method*, *Platform*, *Instrument*), where it can be found (*DatasetDistribution*, *Access*, *Licence*), when it was released and by whom was it created and funded (*Person*, *Organization* and their roles, *Grant*). A set of these DATS core elements also map to the minimal metadata requirements for repositories’ landing pages to support data citation; this work was done by the Repositories Early Adopters Expert Group^12^, part of the Data Citation Implementation Pilot (DCIP) project, an initiative of FORCE11 (https://www.force11.org) and bioCADDIE. The DATS extended elements have been created to progressively accommodate domain-specific metadata for more specialized data types. The current extended set is tailored to DataMed use cases, and therefore specific for life, environmental and biomedical science domains, but it can be further extended as needed.

### Serializations: Schema.org and Bioschemas

Currently the core and the extended DATS entities are available as machine readable JSON schemata, and as a JSON-LD serialization annotated with Schema.org (http://schema.org); serializations in other formats can also be done, if needed. Schema.org is a collaborative, community activity with a mission to create, maintain, and promote schemas for structured data on the Internet, on web pages, in email messages, and beyond. Sponsored by Google, Microsoft, Yahoo and Yandex, Schema.org is already used by over 10 million sites to markup their web pages and email messages. Therefore, anchoring the metadata model to such a potentially powerful vocabulary was one of the early design decisions, outlined in the bioCADDIE white paper. By implementing a Schema.org-annotated DATS model, web-based DataMed will benefit from: an increased visibility by major search engines and tools, increased accessibility via common query interfaces, and possibly, an improvement in search ranking. Conversely, data repositories indexed by bioCADDIE will also get the benefit from being more visible to search engines beyond DataMed.

As Schema.org is a generic vocabulary, its use in the life and biomedical areas requires some extensions. Gaps in coverage have been identified during the annotation process: notifications of these gaps have been submitted to the Schema.org github tracker. The current mapping file, named “Appendix II - NIH BD2K BioCADDIE DataMed DATS v2.1 mapped to schema.org file”, is available as part of the DATS model v2.1 release^10^; this may be subject to change as current Schema.org elements evolve. In addition, members of the bioCADDIE team are working to coordinate extensions to Schema.org for datasets by: participating and leading activities under the Bioschemas umbrella (http://bioschemas.org), also covering and other digital objects; and communicating with the WC3 Healthcore and Schema Vocabulary Community Group (https://www.w3.org/community/schemed), focused towards clinical studies.

## Adoption

Developed iteratively with the input and feedback of a community of international researchers, service providers and knowledge representation experts, DATS is currently implemented by DataMed; this is the intended and primary scope for this metadata model. The bioCADDIE team is going to great lengths to balance a ‘pull’ and ‘push’ approaches to the use of DATS. Initially the effort has been put on the pull approach, requiring the bioCADDIE team to map the target repositories’ model to DATS, to inform the implementation of data harvesting converters, pull data from those repositories and - at the same time - also to test the model in real life and help refine it. Subsequently, the bioCADDIE team has delivered documentation, created examples and held a first workshop with prospective users to lay the basis for the ‘push’ approach, where the owner of the target repository is enabled to export the relevant DATS metadata for indexing in DataMed. To further the uptake, the bioCADDIE team has also released mappings of DATS to a number of existing generic and widely used schemas, such as Schema.org, and domain specific repository schemas. The current mapping file, named “Appendix III - NIH BD2K BioCADDIE DataMed DATS v2.1 mapped to other models file”, is available as part of the DATS model v2.1 release^10^.

Especially notable are our successful collaborations with other data aggregators and service providers that are working to implement DATS. These include - but are not limited to: the Inter-university Consortium for Political and Social Research (ICPSR), the world’s largest archive of digital social science data and one of the NIH-supported repositories (https://www.nlm.nih.gov/NIHbmic/nih_data_sharing_repositories.html); the NIHBD2K OmicsDI^13^, a data discovery index for proteomics, genomics and metabolomics datasets; the NIH Federal Interagency Traumatic Brain Injury Research (FITBIR, https://fitbir.nih.gov), an informatics system to share data across the entire TBI research field; ImmPort (http://www.immport.org), an informatics system supporting the NIH mission to share data, focused on the immunology data; and DataCite (https://www.datacite.org), a global non-profit organisation that supports the creation and allocation of Digital Object Identifiers (DOIs) for research data and accompanying metadata. Another example of use is provided by the NIH BD2K CEDAR metadata authoring tool that works to provide a DATS-compliant template to help researchers to describe and expose their datasets, which are not yet in public repositories, to indexing in DataMed.

To continue these collaborations and foster new ones, driving wider adoption, the bioCADDIE team has also set up a WG on Standards-driven Curation Best Practices (https://biocaddie.org/group/working-group/working-group-12-standard-driven-curation-best-practices) to: (i) assist repositories in creating DATS metadata from the information that they hold; (ii) draw lessons about data curation from experiences of developing data harvesting converters in DataMed; and (iii) encourage best practices that make data FAIR.

## Discussion

A digital ecosystem, such as the NIH Commons, consists of objects (publications and datasets, but also software, code, workflows, training materials, tools, standards, etc.) that are indexed in a consistent fashion, with information related to their origin, contents, and availability. These objects must be findable, accessible, interoperable, and reusable, according to the FAIR principles; this is achieved through a series of technical and social processes involving researchers, informaticians, data scientists, IT professionals, governmental agencies, services providers and publishers.

In this article, we described one portion of the infrastructure, namely the model for the dataset object of this ecosystem. Through a process in which we identified potential use cases and common elements across existing metadata models, we were able to produce a metadata model that is being implemented by the BD2K DataMed prototype, as well as used by other initiatives. The model is expected to continue to evolve with input from the community, and to be used directly by repository managers and data producers to map from their existing metadata. As the model matures, we anticipate new tools and APIs will be developed and the boundary between the shallow indexing intended for DataMed and the deep indexing provided by the repositories and community aggregators to be crossed seamlessly.

## Methods

The DATS model was developed given the following three considerations and approaches.

1. A variety of data discovery initiatives exist or are being developed; although they have different scope, use cases and approaches, the analysis of their metadata schemas is valuable. Several meta-models for representing metadata were reviewed to determine essential items: these have more specific aims and different use cases than the intended capability of DataMed. The reviewed schemas and models are listed in a dedicated BioSharing Collection for bioCADDIE (https://www.biosharing.org/collection/bioCADDIE).
2. Identification of the initial set of metadata elements was based on: (i) analyses of use cases (a *top-down* approach); and (ii) mapping of existing metadata schemas (a *bottom-up* approach), to find convergences and common metadata elements.
3. Use cases have been guiding elements throughout the development process, in order to define the appropriate boundaries and level of granularity. The use cases have been: (i) collected at the bioCADDIE Use Cases Workshop, (ii) extracted from the bioCADDIE White Paper, (iii) provided by the NIH, and (iv) submitted by the community. From these use cases, a set of ‘competency questions’ were derived; these defined the questions that we want DataMed to answer in full, only partially, and which are out of scope. The questions were abstracted, key concepts were highlighted, color-coded and binned in entities, attributes and values categories, to be easily matched with the results of the *bottom-up* approach.

The DATS model v2.1 release^10^ also contains: (i) key examples of competency questions where the concepts highlighted have been binned in entities, attributes and values categories, are provided as part of the “DataMedDATSspecificationv2.1-NIH-BD2KbioCADDIE” file; (ii) a document, named “Appendix I - NIH BD2K BioCADDIE DataMed DATS model v2.1”, detailing the DATS elements that are associated to relevant use cases and/or to the existing schema(s)/model(s) used in the development process, to justify their relevance and provenance; and (iii) the current mapping file, named “Appendix IV - NIH BD2K bioCADDIE WG3 Metadata Mapping File v1.1”, between generic metadata schemas and key life science-specific ones.

## Acknowledgements

This project is funded by grant U24AI117966 from NIAID, NIH as part of the BD2K program. The DATS work in BioSchemas by S-A. S., A. G-B. and P. R-S. is supported by the EU ELIXIR EXCELERATE project (H2020-INFRADEV-1-2015-1 676559). The co-authors are the key investigators and chairs/co-chairs of the DATS development and implementation activities in DataMed. We thank all contributors to the bioCADDIE Consortium and list here the members of the NIH BD2K and of the bioCADDIE Steering Committee: Alison Yao, Dawei Lin, Dianne Babski, George Komatsoulis, Heidi Sofia, Jennie Larkin, Ron Margolis; members of the bioCADDIE’s Descriptive Metadata WG3: Allen Dearry, Carole Goble, Helen Berman, John Westbrook, Kevin Read, Marc Twagirumukiza, Marcelline Harris, Mary Vardigan, Matthew Brush, Melissa Haendel, Michael Braxenthaler, Michael Huerta, Morris Swertz, Rai Winslow, Ram Gouripeddi, Shraddha Thakkar, Weida Tong; and members of the bioCADDIE’s Accessibility Metadata WG7: Alex Kanous, Anne-Marie Tassé, Damon Davis, Frank Manion, Jessica Scott, Kendall Roark, Mark Phillips, Reagan Moore.

## Authors contributions

S-A.S. led the Descriptive Metadata WG3, contributed to the model and its specification and documentation. A.G-B. and P.R-S. co-led on the model development, specification and documentation; A.G-B. developed the DATS validation code and P.R-S. focused on competency questions. G. A. led the Accessibility Metadata WG7 and chaired the use cases workshop, and with J.L. contributed to the model and its specification and documentation. J.G. and H.X. led the implementation of the model and their feedback, along with those from H.K., R.L., Y.L., B.O., X.C., I.F. and A.E.G, contributed to its refinement and releases. S-A.S., A.G-B., P.R-S., J.G., and I.F. worked with the Schema.org and BioSchemas collaborators. L.O-M. led the bioCADDIE consortium and ensured that DATS remained central to the other activities of the NIH Commons ecosystem. She wrote the discussion portion of this manuscript and provided critical edits to other portions of the text. All authors gave iterative feedback on the model and the community engagement and as well as the manuscript. S-A.S., A.G-B. and P.R-S. drafted the first version of the manuscript, with contribution from L.O-M., G.A. and J.G. All coauthors have contributed to its final version.

